# Statistical machine learning of sleep and physical activity phenotypes from sensor data in 96,220 UK Biobank participants

**DOI:** 10.1101/187625

**Authors:** Matthew Willetts, Sven Hollowell, Louis Aslett, Chris Holmes, Aiden Doherty

## Abstract

Current public health guidelines on physical activity and sleep duration are limited by a reliance on subjective self-reported evidence. Using data from simple wrist-worn activity monitors, we developed a tailored machine learning model, using balanced random forests with Hidden Markov Models, to reliably detect a number of activity modes. We show that physical activity and sleep behaviours can be classified with 87% accuracy in 159,504 minutes of recorded free-living behaviours from 132 adults. These trained models can be used to infer fine resolution activity patterns at the population scale in 96,220 participants. For example, we find that men spend more time in both low- and high-intensity behaviours, while women spend more time in mixed behaviours. Walking time is highest in spring and sleep time lowest during the summer. This work opens the possibility of future public health guidelines informed by the health consequences associated with specific, objectively measured, physical activity and sleep behaviours.

## INTRODUCTION

The way that adults spend their time (for example how long they sleep, walk, and sit) has important health implications^1–5^. However this evidence is largely based on self-reported data that are crude and prone to measurement error^6^. Therefore, uncertainty exists on the exact amount and types of sleep and physical activity behaviours that should be recommended, and which interventions and programmes may be most effective in helping people live more healthily. As a result, longitudinal studies now aim to collect objective measures of sleep and physical activity via wrist-worn accelerometers so that their health consequences can be understood^7–10^. As an important first step, ‘vector magnitude’ methods have been developed to objectively measure the volume and intensity levels of physical activity from accelerometer data in large health datasets^10–12^. However, a better understanding of the health consequences of individual lifestyle health behaviours (such as sleeping, sitting, and walking) would arguably help inform public health recommendations that are readily interpretable and implementable. For example, a recommendation of 30 min/day of walking might be more easily understood and actionable than 30 min/day of moderate-to-vigorous intensity physical activity^13^.

Flaws exist in the validation of current methods to extract behavioural information from accelerometer data for relevant biomedical analysis. For example, machine learning methods to detect specific behaviours of interest^14^, such as walking and sitting, have generally not been validated in realistic free-living environments^15^. The validation of these methods in laboratory scenarios is unrealistic as it usually involves a limited number of activities^16^, poor variety within each activity, and an unrealistic relative contribution in time for each activity type^17^. As a result, it is difficult to verify whether current assumptions on the ability to predict walking^18^, or bicycling^15,19^, from sensor data are correct or biased in some manner. Recent work by Ellis and colleagues in a US study has demonstrated that the relevance and accuracy of accelerometer based machine learning methods improves across a range of activities when trained on free-living, rather than controlled laboratory data^20^. However, machine learning methods have not been assessed in large scale health sensor datasets for face validity or investigated to evaluate if they offer behavioural insight.

In this paper we describe the development of a machine learning method to objectively measure lifestyle health behaviours from wrist-worn accelerometer data. We firstly assessed its performance in free-living scenarios using a dataset of 132 adults, 84 of which are female, aged 18-91 who wore an accelerometer and wearable camera (a method comparable to direct-observation^21^). This labelled dataset for machine learning development and held-out validation is many times larger (~159,504 minutes of behaviour) than previous lab-based studies [330 – 3,600 mins^15,18,22^] and free-living studies with short periods of direct observation [3,400 −24,000 mins^23,24^]. We then report the utility of our trained method to assess behavioural variation in more than 100,000 UK Biobank participants aged 43-78 by different self-reported phenotypes. This approach provides an automated analysis of objectively measured behavioural variation in lifestyle behaviours and can be used by researchers to study social and health behaviours at a resolution not previously available.

## RESULTS

**Table 1.**
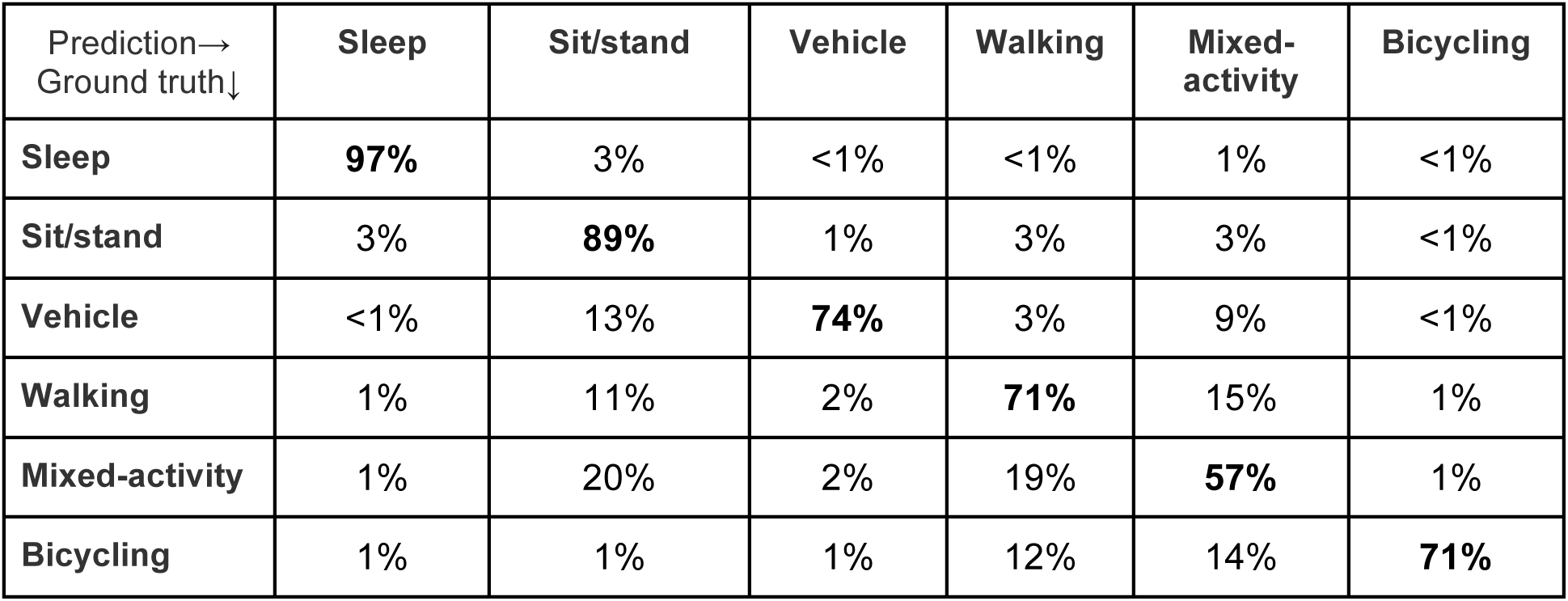
Percentage of machine-learned behaviours automatically classified from wrist-worn accelerometer data. Confusion matrix after leave-one-out validation on 84,616 labelled minutes of human activity in free-living environments: the CAPTURE-24 study 2014-2015 (n = 132).

| Prediction→<br>Ground truth↓ | Sleep | Sit/stand | Vehicle | Walking | Mixed-activity | Bicycling |
| --- | --- | --- | --- | --- | --- | --- |
| Sleep | **97%** | 3% | <1% | <1% | 1% | <1% |
| Sit/stand | 3% | **89%** | 1% | 3% | 3% | <1% |
| Vehicle | <1% | 13% | **74%** | 3% | 9% | <1% |
| Walking | 1% | 11% | 2% | **71%** | 15% | 1% |
| Mixed-activity | 1% | 20% | 2% | 19% | **57%** | 1% |
| Bicycling | 1% | 1% | 1% | 12% | 14% | **71%** |

For activity recognition, we trained a balanced random forest with a Hidden Markov Model containing transitions between predicted activity states and emissions trained using a free-living groundtruth to identify six pre-defined classes of behaviour {bicycling, sit/stand, walking, vehicle, mixed activity, sleep} from accelerometer data. Full details of these models are provided in MATERIALS and METHODS, subsections on activity recognition and time smoothing. For comparison against a free-living groundtruth and to maximise available training data, we conducted leave-one-subject-out cross validation for each of our 132 participants. Over these our model obtained a mean accuracy of 87% with a kappa inter-rater agreement score of 0.81 over all the behaviour types in 30-second windows. Sleep/wake classification was most robust, see our minute-level confusion matrix (Table 1). As expected, there was a wide range of individual variation in classification performance at the daily level, see Bland Altman plots for each activity type (Fig. S1). Overall classification performance was not materially altered by the inclusion of sex as a parameter (Fig. S2). For example, training on all participants from one sex group and then testing on the other, resulted in almost identical overall classifications scores (difference in kappa score <0.0001). Increasing the number of decision trees in the random forest also had little effect on overall classification performance (Fig. S3). Age had a small effect on classification performance, as shown when training on all participants in the top (age>=53) or bottom (age<=29) quartiles and then testing on the other group (kappa = 0.82 trained in old, tested in young; kappa = 0.77 trained in young, tested in old). However, a marked change occurred with the inclusion of hidden Markov model time smoothing, over base random forest predictions, which boosted overall classification performance Kappa score from 0.69 to 0.81 (Table S1). For energy expenditure prediction, we pre-specified 11 classes of behaviour {bicycling, gym, sitstand+activity, sitstand+lowactivity, sitting, sleep, sports, standing, vehicle, walking, walking+activity} that were based on grouping scores in Metabolic Equivalent of Task^25^ (MET). We then took the marginal probability of each state and used this to create a weighted average of the MET scores from the 11 classes. Performing leave-one-subject-out cross validation, we find our model had a root mean squared error of 1.75 MET hours/day (r=0.85). This compares favourably (RMSE of 2.16 MET hours/day and r=0.81) to using random forests for regression^26^ on our dataset.

**Fig. 1.**
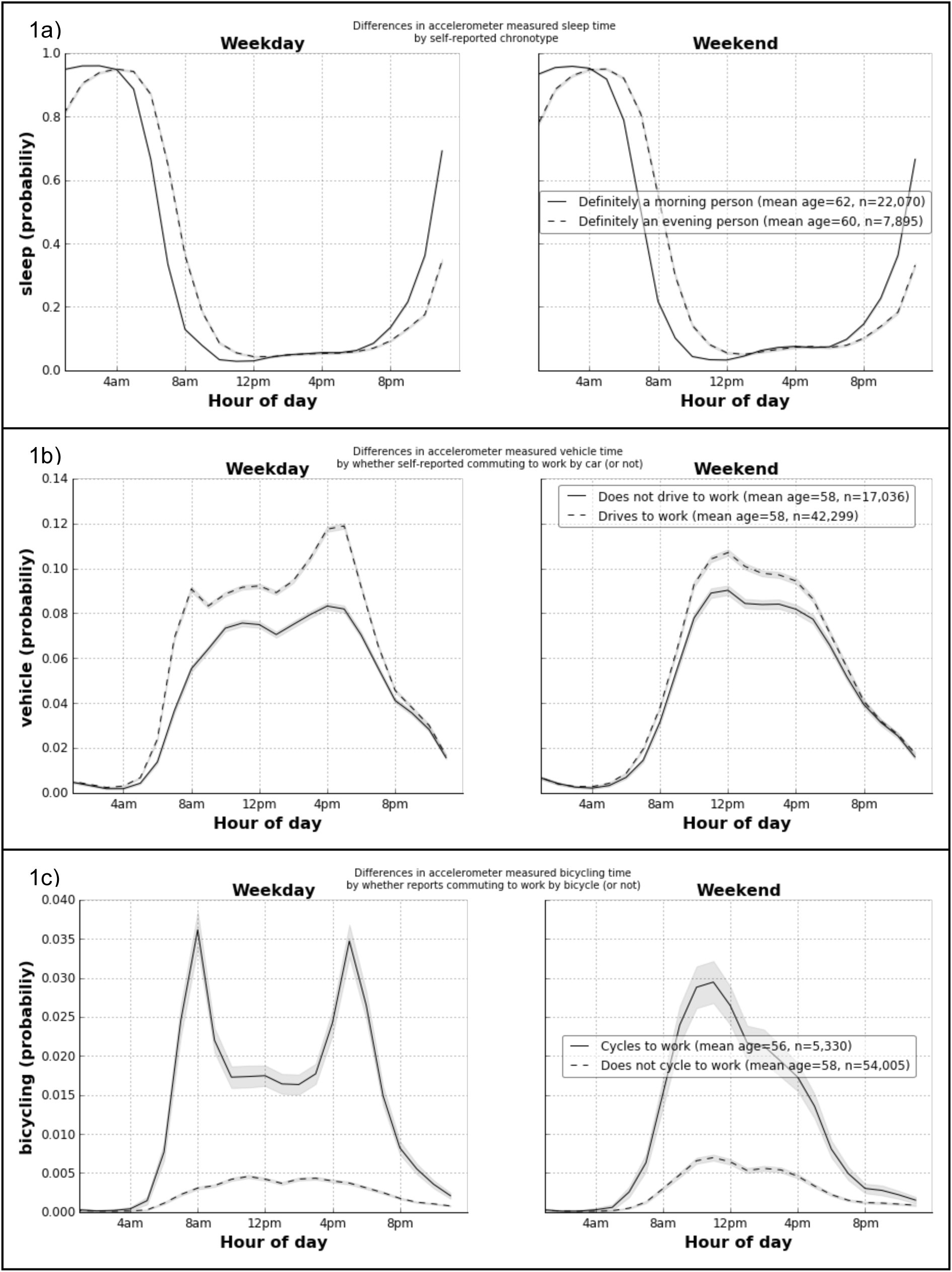
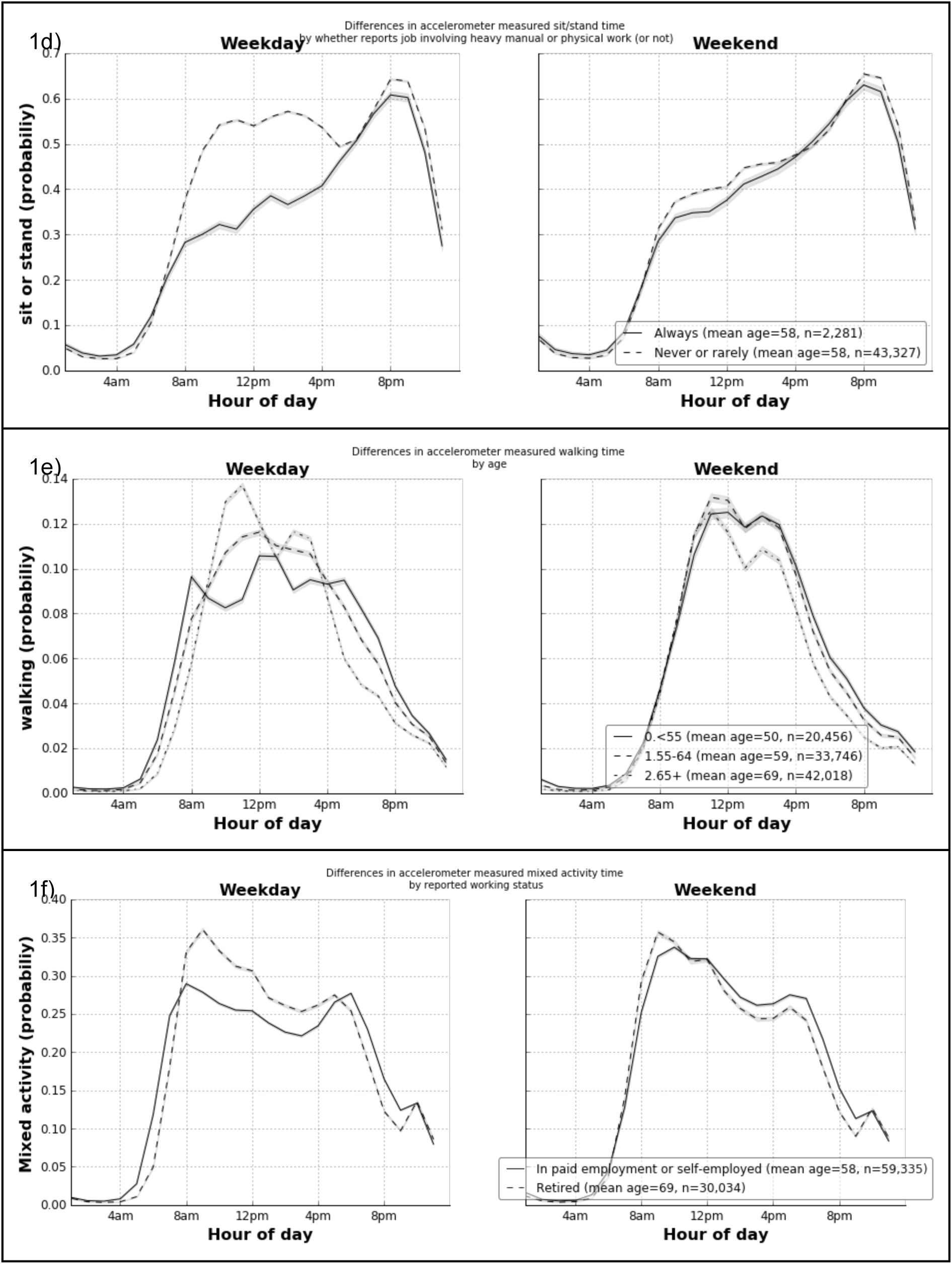
Variation in accelerometer-measured behaviour types across the day by participant characteristics (measured 2007-2010) and weekday/weekend (2013-2015): the UK Biobank study (n = 96,220).

To assess the face validity of our activity recognition method in a large prospective dataset, we applied our model to 103,712 UK Biobank participants. We removed participants who did not wear the device for a sufficient amount of time (n=7,128), or who had device errors^10^ (n=364). On 96,220 participants, we then plotted each aggregated activity by time-of-day and stratified groups by characteristics self-reported during the study baseline visit a mean of 5.7 years before accelerometer wear (see Fig. 1). Fig. 1a shows self-reported ‘evening’ people were more likely to be classified as sleeping at 8am on weekends vs. ‘morning’ people (55% vs. 22%, p<10^−100^ after adjustment for age, sex, ethnicity, area deprivation, smoking, alcohol, fruit/veg intake, and self-rated overall health). Self-reported car users were more likely to be classified as driving at 8am on weekdays (9.1% vs. 5.5%, p<10^−100^ after adjustment for other factors) (see Fig. 1b). Similarly, self-reported cyclists were more likely to be classified as cycling at 8am on weekdays (3.6% vs. 0.3%, p<10^−100^ after adjustment for other factors). Those in active occupations were more likely not to be classified as sitting or standing at 11am on weekdays than office based workers (55% vs. 31%, p<10^−100^ after adjustment for other factors) (Fig. 1d). Older adults (aged 65+) were more likely to be classified as walking at 11am on weekdays than younger adults (aged <55) (14% vs. 9%, p=3×10^−18^ after adjustment) (Fig. 1e). Finally, retired people were more likely to be classified as doing mixed activity at 11am on weekdays than their working counterparts (31% vs. 26%, p=1.3×10^−38^ after adjustment). As expected, these age/occupation differences mostly disappear on weekends (Fig. 1d-f).

**Table 2.** Objective machine-learned measures of physical activity (vector magnitude), sleep, walking, sitting-or-standing, bicycling, vehicle, and mixed activity time: the UK Biobank study 2013-2015 (n = 96,220).

|  | Individuals | Physical activity | MET | Sleep | Walking | Sit / stand | Bicycling | Vehicle | Mixed |
| --- | --- | --- | --- | --- | --- | --- | --- | --- | --- |
|  | [ n ] | [mg] | [MET hrs/day] |  |  |  |  |  |  |
| Age |  |  |  |  | [% time] |  |  |  |  |
|  |  |  |  |  | [mean ± stdev] |  |  |  |  |
| <55 | 20,456 | 31.7 ± 9.1 | 37.6 ± 3.1 | 36.2 ± 5.0 | 5.5 ± 3.0 | 35.7 ± 7.8 | 0.4 ± 1.0 | 5.3 ± 3.4 | 17.0 ± 7.2 |
| 55-64 | 33,746 | 29.4 ± 8.2 | 37.1 ± 3.0 | 36.7 ± 5.1 | 5.5 ± 3.0 | 35.4 ± 7.6 | 0.3 ± 0.8 | 5.1 ± 3.2 | 17.1 ± 6.9 |
| 65+ | 42,018 | 26.3 ± 7.3 | 36.3 ± 2.9 | 37.3 ± 5.4 | 5.1 ± 3.0 | 36.1 ± 7.5 | 0.2 ± 0.6 | 4.5 ± 2.8 | 16.9 ± 6.7 |
| p value<sup>A</sup> |  | <1x10<sup>-300</sup> | <1x10<sup>-300</sup> | 5x10<sup>-148</sup> | 3x10<sup>-93</sup> | 2x10<sup>-31</sup> | 2x10<sup>-146</sup> | 7x10<sup>-270</sup> | 2x10<sup>-04</sup> |
| Cohen's d |  | .66 | .43 | .21 | .13 | .08 | .20 | .27 | .03 |
| **Sex** |  |  |  |  |  |  |  |  |  |
| Women | 54,158 | 29.0 ± 8.0 | 37.1 ± 2.9 | 36.9 ± 4.9 | 4.8 ± 2.7 | 34.6 ± 7.2 | 0.2 ± 0.6 | 4.6 ± 2.8 | 18.9 ± 6.7 |
| Men | 42,062 | 28.0 ± 8.7 | 36.5 ± 3.2 | 36.7 ± 5.6 | 5.9 ± 3.3 | 37.3 ± 7.8 | 0.4 ± 1.0 | 5.2 ± 3.4 | 14.5 ± 6.4 |
| p value<sup>A</sup> |  | 5x10<sup>-33</sup> | 1x10<sup>-197</sup> | 3x10<sup>-17</sup> | <1x10<sup>-300</sup> | <1x10<sup>-300</sup> | <1x10<sup>-300</sup> | 1x10<sup>-248</sup> | <1x10<sup>-300</sup> |
| Cohen's d |  | .11 | .21 | .04 | .37 | .37 | .25 | .20 | .68 |
| **Self-rated health** |  |  |  |  |  |  |  |  |  |
| Excellent | 21,101 | 30.8 ± 8.9 | 37.4 ± 2.9 | 36.5 ± 4.8 | 5.6 ± 3.0 | 35.0 ± 7.2 | 0.4 ± 1.0 | 5.0 ± 3.0 | 17.5 ± 6.8 |
| Good | 57,792 | 28.6 ± 8.0 | 36.9 ± 3.0 | 36.8 ± 5.1 | 5.3 ± 3.0 | 35.5 ± 7.4 | 0.3 ± 0.7 | 4.9 ± 3.1 | 17.2 ± 6.8 |
| Fair | 15,313 | 26.1 ± 7.8 | 36.0 ± 3.1 | 37.2 ± 5.8 | 4.9 ± 3.1 | 37.1 ± 8.2 | 0.2 ± 0.6 | 4.7 ± 3.3 | 15.9 ± 7.1 |
| Poor | 2,707 | 23.3 ± 7.8 | 34.9 ± 3.5 | 37.9 ± 7.1 | 3.8 ± 3.0 | 39.2 ± 9.1 | 0.2 ± 0.6 | 4.2 ± 3.2 | 14.6 ± 7.2 |
| p value<sup>A</sup> |  | <1x10<sup>-300</sup> | <1x10<sup>-300</sup> | 8x10<sup>-54</sup> | 2x10<sup>-271</sup> | 3x10<sup>-193</sup> | 6x10<sup>-149</sup> | 1x10<sup>-33</sup> | 3x10<sup>-106</sup> |
| Cohen's d |  | .90 | .79 | .24 | .60 | .51 | .27 | .26 | .40 |
| **Time of day** |  |  |  |  |  |  |  |  |  |
| 0-5.59<sub>am</sub> | 96,220 | 5.0 ± 3.7 | 23.9 ± 1.9 | 92.3 ± 10.0 | 0.3 ± 1.0 | 5.7 ± 7.8 | 0.0 ± 0.2 | 0.4 ± 1.8 | 1.3 ± 2.7 |
| 6-11.59<sub>am</sub> | 96,220 | 38.8 ± 15.4 | 41.3 ± 5.5 | 29.1 ± 13.5 | 7.2 ± 5.5 | 32.7 ± 11.8 | 0.4 ± 1.4 | 5.9 ± 4.8 | 24.6 ± 11.0 |
| 12-5.59<sub>pm</sub> | 96,220 | 44.4 ± 14.9 | 45.7 ± 5.4 | 4.7 ± 6.3 | 10.1 ± 6.3 | 49.6 ± 13.4 | 0.5 ± 1.6 | 9.1 ± 6.1 | 26.0 ± 12.9 |
| 6-11.59<sub>pm</sub> | 96,220 | 26.2 ± 10.9 | 36.3 ± 4.8 | 21.2 ± 13.0 | 3.5 ± 3.3 | 55.0 ± 12.9 | 0.2 ± 0.8 | 4.1 ± 4.4 | 16.0 ± 8.7 |
| p value<sup>B</sup> |  | <1x10<sup>-300</sup> | <1x10<sup>-300</sup> | <1x10<sup>-300</sup> | <1x10<sup>-300</sup> | <1x10<sup>-300</sup> | <1x10<sup>-300</sup> | <1x10<sup>-300</sup> | <1x10<sup>-300</sup> |
| Cohen's d |  | 3.6 | 5.4 | 10.5 | 2.2 | 4.6 | .44 | 1.9 | 2.6 |
| **Day** |  |  |  |  |  |  |  |  |  |
| Weekday | 96,220 | 28.7 ± 8.5 | 37.0 ± 3.2 | 36.2 ± 5.5 | 5.4 ± 3.2 | 36.2 ± 8.2 | 0.3 ± 0.8 | 5.0 ± 3.4 | 16.9 ± 7.3 |
| Weekend | 96,220 | 28.2 ± 9.9 | 36.4 ± 3.6 | 38.3 ± 6.6 | 5.1 ± 3.7 | 34.7 ± 8.7 | 0.3 ± 1.1 | 4.5 ± 3.9 | 17.1 ± 7.8 |
| p value<sup>B</sup> |  | 4x10<sup>-98</sup> | <1x10<sup>-300</sup> | <1x10<sup>-300</sup> | 4x10<sup>-172</sup> | <1x10<sup>-300</sup> | .765 | <1x10<sup>-300</sup> | 1x10<sup>-27</sup> |
| Cohen's d |  | .05 | .17 | .35 | .09 | .17 | .00 | .15 | .03 |
| **Season** |  |  |  |  |  |  |  |  |  |
| Spring | 21,839 | 29.0 ± 8.5 | 37.0 ± 3.0 | 36.7 ± 5.1 | 5.4 ± 3.1 | 35.6 ± 7.6 | 0.3 ± 0.8 | 4.9 ± 3.1 | 17.0 ± 6.9 |
| Summer | 25,273 | 29.1 ± 8.5 | 37.0 ± 3.0 | 36.3 ± 5.1 | 5.3 ± 3.1 | 35.7 ± 7.7 | 0.3 ± 0.9 | 5.0 ± 3.1 | 17.4 ± 7.0 |
| Autumn | 28,699 | 28.4 ± 8.2 | 36.8 ± 3.0 | 36.8 ± 5.2 | 5.3 ± 3.0 | 35.9 ± 7.5 | 0.3 ± 0.7 | 4.9 ± 3.1 | 16.8 ± 6.8 |
| Winter | 20,409 | 27.6 ± 8.0 | 36.5 ± 3.0 | 37.5 ± 5.4 | 5.0 ± 2.9 | 35.9 ± 7.5 | 0.2 ± 0.6 | 4.7 ± 3.0 | 16.7 ± 6.8 |
| p value<sup>A</sup> |  | 6x10<sup>-107</sup> | 3x10<sup>-72</sup> | 3x10<sup>-136</sup> | 5x10<sup>-46</sup> | 4x10<sup>-05</sup> | 1x10<sup>-77</sup> | 2x10<sup>-32</sup> | 2x10<sup>-32</sup> |
| Cohen's d |  | .18 | .15 | .23 | .14 | .04 | .16 | .11 | .10 |

<sup>A</sup> Age, sex, self-rated health, season (Spring starting on 1<sup>st</sup> March): Two-way analysis of variance test used to compare metrics between groups adjusting for age, sex, ethnicity, area-deprivation, smoking, alcohol, self-rated health and season of wear.
<sup>B</sup> Time of day, day: Repeated two-way analysis of variance test used to compare metrics within individuals and between groups adjusting for age, sex, ethnicity, area-deprivation, smoking, alcohol, self-rated health, and season of wear.

Table 2 describes the variation in accelerometer-measured total time for each behaviour type, by age, self-rated health, time-of-day, weekday/weekend, and season. Younger participants spent more time in active behaviours than their older counterparts (e.g. 5.5% vs. 5.1% time for walking, p=3×10^−93^). However, these age differences for specific behaviours weren’t as pronounced as for vector magnitude, which is a proxy measure of overall physical activity^10^. The behaviour patterns of men appeared more polarised than that of women, with more time in low-intensity activities such as sitting (37.3% vs. 34.6%, p<10^−100^) but also more time in purposeful activity behaviours such as walking (5.9% vs. 4.8%, p<10^−100^) and bicycling. Women spent more time engaged in mixed activity behaviours than men (18.9% vs. 14.5%, p<10^−100^). Self-rated health differences were strongest for traditional vector magnitude measures, but also noticeable for walking time between those in excellent versus poor self-rated health (5.6% vs. 3.8%, p<10^−100^). While overall physical activity differences between weekdays and weekends were small (Cohen’s d=0.05), behavioural differences were more noticeable (d=0.35 and d=0.17 for longer sleep and less sitting time at weekends respectively). Small seasonal differences also existed with walking time highest in spring, and in summer versus winter there was less sleep (36.3% vs. 37.5%, p<10^−100^), more bicycling (0.3% vs. 0.2%, p=1×10^−77^), and higher energy expenditure (37.0 vs. 36.5 MET hours/day, p=3×10^−72^).

**Fig. 2.**
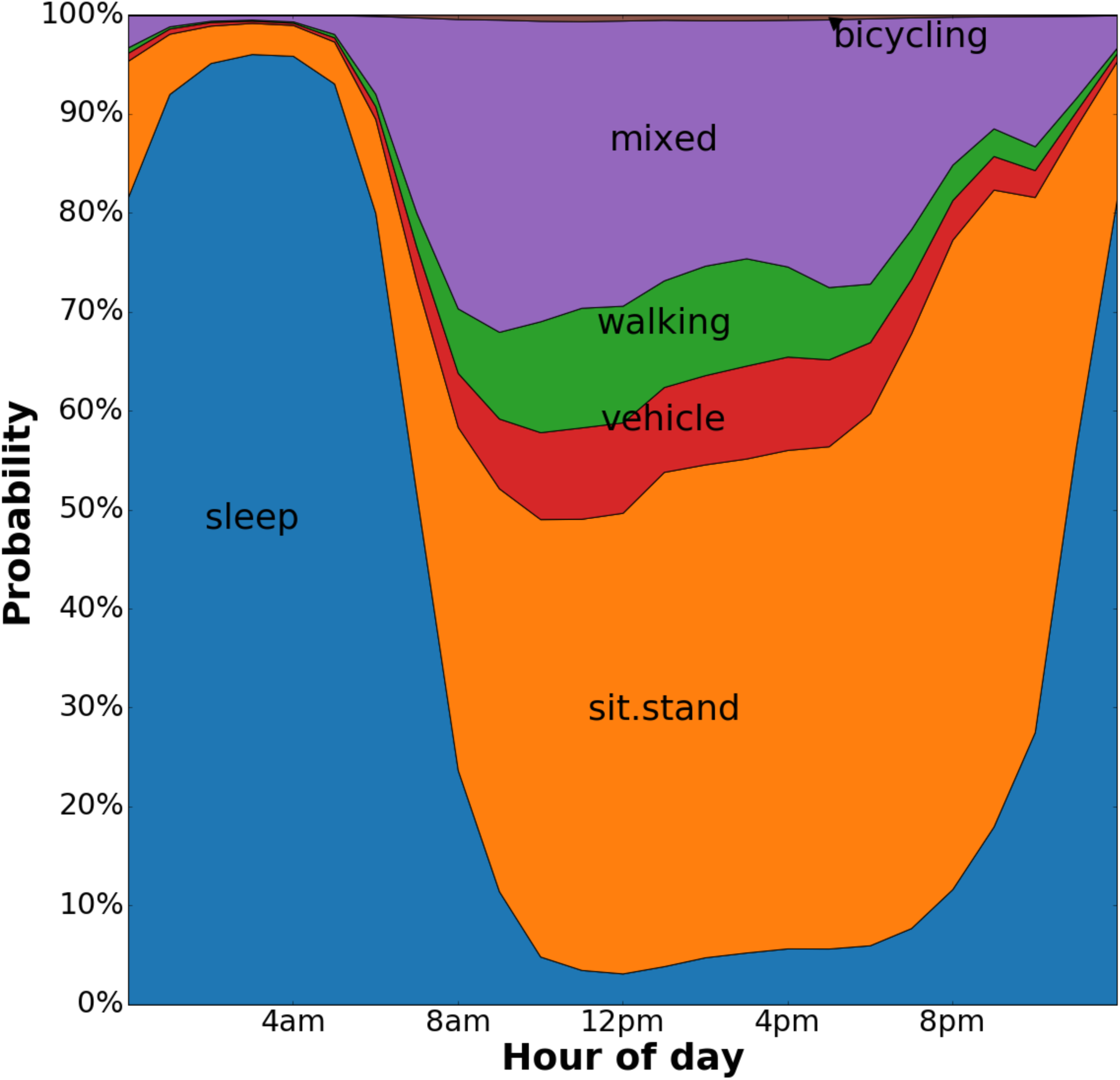
Variation in accelerometer-measured time by activity type: the UK Biobank study 2013-2015 (n = 96,220).

Due to the time-series nature of the data, it is possible to illustrate the likelihood for each activity state throughout the day in 96,220 UK Biobank participants (see Fig. 2), and how this relates to daily energy expenditure (see Fig. S4). Apart from mixed-activity and METs (r=0.75), the different activity types are weakly correlated (absolute overall mean r=0.37), indicating their utility as new sources of information (Fig. S5).

## DISCUSSION

This study represents the largest ever assessment of objectively measured sleep and physical activity behaviours using state-of-the-art machine learning methods trained with a free-living groundtruth. To our knowledge, this is the first time that {“bicycling”, “mixed”, “sit/stand”, “sleep”, “vehicle”, “walking”} behaviours have been objectively measured in a large-scale dataset. We have demonstrated the feasibility of our method as scale in 96,220 UK Biobank participants where, for example, commute times can be seen for those who self-report as cycling to work. The objective and fine-grained measures (with ~20,000 behavioural predictions per person per week) that we have developed will help more precisely understand the effectiveness of treatments and also the disease processes associated with behaviour variation.

The overall classification score of kappa=0.81 for our method (a balanced random-forest with Markov transitions on predicted states and emissions trained in a naturalistic free-living scenario) represents a substantial level of agreement with the wearable camera groundtruth^27^. This is comparable to the level of performance (kappa=0.80) that we expect from humans annotating behaviour from wearable camera data^28,29^. Our overall crude accuracy of 87% is at least as good as reported in other free-living studies with wrist-worn accelerometers [85% in^30^ and 61% in^23^]. Direct comparison across such studies is difficult due to heterogeneity across populations, devices, and definition of behaviour labels. For energy expenditure prediction, random forests for regression^26^ perform better than linear models for wrist-worn data^22^, and we found the inclusion of activity labels reduced noise and thus further improved performance (e.g. movement is high when in a vehicle, but energy expenditure is low). This mirrors findings from studies that used older hip-worn accelerometers^31^.

Our study shows that bicycling can be reliably detected from accelerometer data, an activity previously difficult to classify in laboratory studies^15,19^. Potential explanations for this might be our use of devices with higher sampling rates (100Hz vs. 1Hz)^19^ that can capture important bicycling activities, or models developed in free-living moving bicycles rather than stationary laboratory bicycles^15^. Previous laboratory studies indicated that walking can be classified with a high degree of accuracy (sensitivity and specificity both >90%). However, our data and that from Ellis et al.^30^ shows that walking is challenging to classify in free-living conditions (sensitivity=0.71, specificity=0.96), probably due to it being part of many everyday activities.

While the participants in our free-living dataset are not a random subsample of the UK Biobank study, the size of our training dataset has helped provide a diverse set of representative behaviours, rather than individuals, which is important for model development. The only other study comparable in training set size used a similar study procedure^30^, but in a US population subgroup and thus could not be reliably extrapolated to UK Biobank data. Our study also uses more stringent evaluation criteria (kappa versus balanced accuracy) that consider unbalanced free-living data where infrequent behaviours are more susceptible to misclassification. Our method is device agnostic, and could be reused in other large sensor datasets^10,11,32,33^, provided model tuning takes place in a relevant population with free-living groundtruth validation tools such as wearable cameras^13,30^. For this study we did not use traditional cumbersome methods to collect sleep^34^ and energy expenditure^35^ groundtruth data, as we preferred to use proxy reference methods for free-living assessment at scale^35,36^. We have not generalised the overall descriptive findings to the UK population since the UK Biobank was established as an aetiological study rather than one aimed at population surveillance^9,37^.

In summary, we describe the first application, to our knowledge, of machine learning to objectively measure lifestyle health behaviours from sensor data in a large prospective health study. Our method has demonstrated substantial agreement with a free-living groundtruth, and shows face validity in a large health dataset. It is now possible to study the sociological and health consequences of behaviour variation in unprecedented detail. The summary variables that we have constructed are now part of the UK Biobank dataset and can be used by researchers as exposures, confounding factors or outcome variables in future health analyses.

## METHODS

### Participants

For the development and free-living evaluation of accelerometer machine learning methods, 143 participants were recruited to the CAPTURE-24 study where adults aged 18-91 were recruited from the Oxford region in 2014-2015^38^. Participants were asked to wear a wrist-worn accelerometer for a 24-hour period and then given a £20 voucher for taking part in this study that received ethical approval from University of Oxford (Inter-Divisional Research Ethics Committee (IDREC) reference number: SSD/CUREC1A/13-262). We removed 11 participants who had missing camera or accelerometer data, or where both sources could not be time-aligned, leaving 132 participants for classifier development. For extrapolation to a large health dataset, we used the UK Biobank dataset where 103,712 participants agreed to wear a wrist-worn accelerometer for a seven day period between 2013-2015^10^. UK Biobank is a large prospective study of 500,000 participants that has collected, and continues to collect, extensive phenotypic and genotypic details about its participants, with ongoing longitudinal follow-up for a wide range of health-related outcomes^37^. Demographic and behavioural variables were recorded by a self-completed touchscreen questionnaire during clinic visits between 2006-2010 (see appendix 1). This study (UK Biobank project #9126) was covered by the general ethical approval for UK Biobank studies from the NHS National Research Ethics Service on 17th June 2011 (Ref 11/NW/0382). As per informed consent procedures, informed consent was obtained and all participant data was anonymised. Methods reported in this manuscript were performed in accordance with relevant guidelines and regulations covered by the aforementioned ethics approval committees.

### Accelerometer

Participants in both studies were asked to wear an Axivity AX3 wrist-worn triaxial accelerometer on their dominant hand at all times. It was set to capture tri-axial acceleration data at 100Hz with a dynamic range of +-8g. This device has demonstrated equivalent signal vector magnitude output on multi-axis shaking tests^39^ to the GENEActiv accelerometer which has been validated using both standard laboratory and free-living energy expenditure assessment methods^36,40^.

### Groundtruth

To construct a groundtruth of reference behaviours, participants in the Oxford study were asked to wear a Vicon Autographer wearable camera while awake on the study measurement day. Wearable cameras automatically take photographs every ~20 seconds, have up to 16 hours battery life and storage capacity for over one week’s worth of images^41^. When worn, the camera is reasonably close to the wearer’s eye line and has a wide-angle lens to capture everything within the wearer’s view^42^. Each image is time-stamped so duration of active travel^43^, sedentary behaviour^29^, and a range of other physical activity behaviours^44^ can be captured. Camera data strongly agrees with more expensive direct observation methods to classify activity types [kappa=0.92^21^]. We used specific ethical guidance for wearable camera research to inform the development of protocols^45^. Images were annotated by human annotators using codes from the compendium of physical activities^25^, using specific wearable camera browsing software^46^ (Doc S1). For quality control, our annotators firstly had to achieve a kappa inter-rater agreement score of >0.8 on separate training data. To extract sleep information, participants were asked to complete a simple sleep diary, as used in the Whitehall study, which consisted of two questions^47^: *‘what time did you first fall asleep last night?’* and *‘what time did you wake up today (eyes open, ready to get up) ?’*. Participants were also asked to complete a HETUS time-use diary^48^, and sleep information from here was extracted in cases where data was missing from the simple sleep diary. This multi-instrument groundtruth resulted in 213 activity labels which were then condensed into six free-living behaviour labels {“bicycling”, “mixed”, “sit/stand”, “sleep”, “vehicle”, “walking”} (see mappings at appendix 2a). Fig. S5 shows a visual representation of the structure and time-balance of labels annotated from this free-living dataset. For energy expenditure metabolic equivalent of task (MET) prediction, we used eleven behaviour labels {“bicycling”, “gym”, “sitstand+activity”, “sitstand+lowactivity”, “sitting”, “sleep”, “sports”, “standing”, “vehicle”, “walking”, “walking+activity”} (see mappings at appendix 2b), each with an associated Metabolic Equivalent of Task (MET) score from the compendium of physical activities.

### Accelerometer Data Preparation

For data pre-processing we followed procedures used by the UK Biobank accelerometer data processing expert group^10^, that included device calibration^49^, resampling to 100Hz, and removal of noise and gravity^10,32,33^. For every non-overlapping 30-second time window, which corresponds to the granularity of groundtruth labels, we then extracted a 126-dimensional feature vector. Our features are listed in Fig. S2 and were selected from an extensive list of time and frequency domain features described in other studies^15,18,30,50^. These included: euclidean norm minus one with negative values truncated to zero^10^, it’s mean, standard deviation, coefficient of variation, median, min, max, 25^th^ & 75^th^ percentiles, mean amplitude deviation, mean power deviation, kurtosis & skew, and Fast Fourier Transform (FFT) 1-15Hz. Features also included the following in each axis of movement: mean, range, standard deviation, covariance, and FFT 1-15Hz. Roll, pitch, yaw, x/y/z correlations, frequency and power bands were also extracted.

### Activity Classification

For activity classification we use random forests^51^ which offer a powerful nonparametric discriminative method for multi-class classification that offers state-of-the-art performance^52^. Predictions of a random forest are an aggregate of individual CART trees (Classification And Regression Trees). CART trees are binary trees consisting of split nodes and terminal leaf nodes. In our case, each tree is constructed from a training set of feature data along with ground truth activity classes. For a standard random forest, to train a tree from *N* data points with *F* features, we first select *N* data points with replacement and 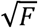 feature variables (without replacement)^51^, then carry out the CART algorithm (Appendix S3). To run each tree for a new data point we follow the decision process of that CART tree where the output is a unit vote for an activity class. One can describe a single tree as a function of a data point *x* that returns a one-hot vector vote for a given class *k*:

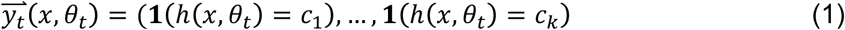

where {*θ_i_*} in equation (1) is the set of parameters describing what thresholds have been chosen and which data points and features have been used in that tree. *h*(·,·) outputs the predicted value of class which is transformed into a one-hot encoding by **1**(·) the indicator function.

The trees individually have high variance so their votes are combined together. This is called ‘bagging’ (from bootstrap aggregating). The combination of trees forms a random forest^51^ of *T* trees given in equation (2):

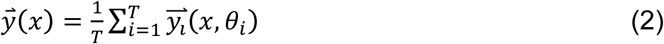

We can either simply take the most commonly voted class as the prediction, or as in equation (3) normalise the votes by the number of trees to get probabilities:

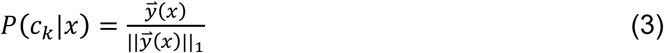

There is randomness in {*θ_i_*}, as we only give each tree a subset of data and features. This ensures that the trees have low correlation and is necessary as the CART algorithm itself is deterministic. Given the unbalanced nature of our dataset, where some behaviours occur rarely, we use balanced Random Forests^53^ to train each tree with a balanced subset of training data. If we have n_rare_ instances of the rarest class, we pick n_rare_ samples, with replacement, of data of each of our classes to form our training set for each tree. As each tree is given only a small fraction of data, we make many more trees than in a standard random forest so that the same number of data points are sampled in training as with a standard application of random forests. We evaluated different numbers of trees in our random forest, each trained using n_rare_ datapoints from each class of activity (Fig. S6).

### Time Smoothing

Random forests are able to classify datapoints, but do not have an understanding of our data as having come from a time series. Therefore we use a hidden Markov model^54^ (HMM) to encode the temporal structure of the sequence of classes and thus obtain a more accurate sequence of predicted classes. A hidden Markov model is a state space model consisting of a sequence of hidden discrete states There is a stochastic sequence of states *ẕ* = {*z*_1_, *z*_2_…, *z*_*t*−1_, *z_t_*, *z*_*t*+1_, …} that have the Markov property that only the present influences the future:

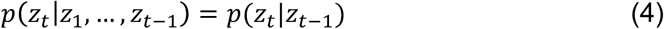

At each time step *t in* *equation* (4), *z_t_* can take one of *k* classes {*c_1_, c_2_, …, c_k_*} and thus the dynamics are described by the transition matrix *k* by *k* in equation (5):

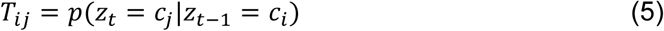

Although we do not observe the hidden states, at each time step there is an observed, stochastic emission *y_t_* that depends on the hidden state *z_t_*. They are drawn from a probability distribution *p*(*y_t_*|*z_t_, ϕ*) where *ϕ* are the various parameters that describe the distribution. They form a sequence, as outlined in equation (6):

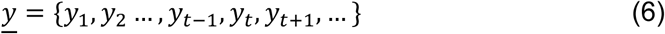

For us the hidden state space sequence *ẕ* of the HMM is the sequence of true activities and the emissions 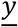 are the predicted activities from the balanced random forest (Fig S7). We thus wish to use our imperfect, noisy, predictions of activity from our random forest to infer the most likely sequence of true activity states that would have given rise to those random forest predictions. The transition matrix and emission distribution were empirically calculated.

The transition matrix and emission distribution are empirically calculated. The transition matrix is calculated from the training set sequence of activity states. The calculation of emission probabilities comes from the out of bag class votes of the random forest. Recall that in a random forest each tree is trained on a subset of the training data. Thus by passing through each tree the training data that it was not trained on we get an estimate of the error of the forest. This gives us directly the probability of predicting each class given the true activity class, which is the emission distribution *p*(*y_t_*|*z_t_, ϕ*) we need. And so the confidence of the random forest in the accuracy of its predictions for an activity follows through into how confident the HMM is that a random forest prediction corresponds to the true activity classification. With this empirically defined HMM, we can then run the Viterbi algorithm^55^ to find the most likely sequence of states *ẕ\** given a sequence of observed emissions in equation (7):

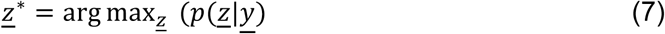

This smoothing corrects erroneous predictions from the random forest, such as where the error is a blip of one activity surrounded by another and the transitions between those two classes of activity are rare. The overall most likely state sequence *ẕ\** is not the same as the sequence of marginally most probable states - for instance there could be forbidden transitions between two sequentially marginally most probable states, rendering that sequence impossible. This is relevant for us as some transitions do not appear in our data set.

### MET Prediction

To predict the MET score we follow the same process of feature extraction, random forest training and HMM definition, but for the eleven-class MET-relevant behaviour labels. However, instead of selecting the Viterbi path we obtain the sequence of marginal probabilities for being in each state at each time given the sequence of observations. Each of the eleven classes of behaviour is a mix of different activities from the compendium of physical activities, thus a representative MET score was calculated by taking the mean of the MET scores used to construct that class from the training dataset. Finally, the predicted MET score for each 30-second chunk is calculated as the assigned MET scores for each of the 11 states, weighted by the marginal probabilities of being in each of those states.

### Extrapolation to large health datasets

We trained a model using all free-living groundtruth data, and applied it to predict behaviour for each 30-second epoch in 103,712 UK Biobank participants’ accelerometer data. For any given time window (e.g. one hour, one day, etc.) the probability of a participant engaging in a specific behaviour type was expressed as the number-of-epoch-predictions-for-class divided by the number-of-epochs. Device non-wear time was automatically identified as consecutive stationary episodes lasting for at least 60 minutes^10^. These non-wear segments of data were imputed with the average of similar time-of-day data points, for each behaviour prediction, from different days of the measurement. We excluded participants whose data could not be calibrated, had too many clipped values^10^, had unrealistically high values (average vector magnitude > 100mg), or who had poor wear-time. We defined minimum wear time criteria as having at least three days (72 hours) of data and also data in each one-hour period of the 24-hour cycle^10^.

### Statistical Analysis

To compare machine predicted behaviour from accelerometer data against the free-living groundtruth, we used leave-one-subject-out cross validation, and reported kappa scores for agreement (unit = 30 second time windows). The Kappa test reflects the inter-rater agreement between two sources taking into account the likelihood of them agreeing by chance^56^. We used Bland-Altman plots to illustrate daily-level summary agreement between predicted behaviour and the groundtruth. For the UK Biobank dataset, descriptive statistics were used to report accelerometer measured time in {“bicycling”, “mixed”, “sit/stand”, “sleep”, “vehicle”, “walking”} behaviours. Age groups were categorised into <55, 55-64, and 65+ years. To quantify statistical differences by age, sex, self-rated health, and season, two-way ANOVA linear regression were used, with analysis adjusted for age, sex, ethnicity, area-deprivation, smoking, alcohol, and self-rated-health. Time-of-day (six hour quadrants) and weekday vs. weekend differences in behaviour were reported using two-way repeated measures ANOVA. As this is used for quantification, rather than hypothesis testing, we report p-values uncorrected for multiple testing. 24-hour activity plots stratified by weekend were used to illustrate accelerometer-measured behavioural profiles. Stack charts were plotted to illustrate the distribution of all objectively measured behavioural types in UK Biobank participants. We used R to perform all statistical analyses^57^.

## Data and code availability

Upon publication, the summary variables that we have constructed will be made available as a part of the UK Biobank dataset at http://biobank.ctsu.ox.ac.uk/crystal/label.cgi?id=1008. All data processing, feature extraction, machine learning, and analysis code will be available at https://github.com/activityMonitoring.

## ACKNOWLEDGEMENTS

We would like to thank all participants for agreeing to volunteer in this research. For data collection at Oxford and annotation we would like to acknowledge the support of Jonathan Gershuny, the UK Economic and Social Research Council (grant number ES/L011662/1), Charlie Foster, Paul Kelly, Teresa Harms, Emma Thomas, Karen Milton, Wong Tsz Yan, Nicole Gray, and Salma Haque. We would like to thank Martin Landray for feedback on this work. The UK Biobank Activity Project and the collection of activity data from participants was funded by the Wellcome Trust (https://wellcome.ac.uk/) and the Medical Research Council (http://www.mrc.ac.uk/). The analysis was supported by the NIHR Biomedical Research Centre, Oxford [AD]; the British Heart Foundation Centre of Research Excellence at Oxford (http://www.cardioscience.ox.ac.uk/bhf-centre-of-research-excellence) [grant number RE/13/1/30181 to AD]; the Li Ka Shing Foundation (http://www.lksf.org/) [to AD]; and the Engineering and Physical Sciences Research Council (EPSRC) [grant number EP/G03706X/1 for MW]. We would also like to acknowledge the use of the University of Oxford Advanced Research Computing (ARC) facility in carrying out this work. http://dx.doi.org/10.5281/zenodo.22558 The MRC and Wellcome Trust played a key role in the decision to establish UK Biobank, and the accelerometer data collection. No funding bodies had any role in the analysis, decision to publish, or preparation of the manuscript.

## AUTHOR CONTRIBUTIONS STATEMENT

A.D., C.H., and L.A. conceived this study. M.W., S.H., and A.D. developed and implemented the statistical models. S.H. and A.D. developed and implemented the population inference models. A.D., M.W., and S.H. wrote the paper. A.D., C.H., L.A., M.W., and S.H. were involved in interpreting results and editing the manuscript.

## ADDITIONAL INFORMATION

The authors declare no competing interests.

